# Genome-scale quantification and prediction of pathogenic stop codon readthrough by small molecules

**DOI:** 10.1101/2023.08.07.552350

**Authors:** Ignasi Toledano, Fran Supek, Ben Lehner

## Abstract

Premature termination codons (PTCs) cause ∼10-20% of Mendelian diseases and are the major mechanism of tumor suppressor gene inactivation in cancer. A general strategy to alleviate the effects of PTCs would be to promote translational readthrough. Nonsense suppression by small molecules has proven effective in diverse disease models, but translation into the clinic is hampered by ineffective readthrough of many PTCs. Here we directly tackle the challenge of defining drug efficacy by quantifying readthrough of ∼5,800 human pathogenic stop codons by 8 drugs. We find that different drugs promote readthrough of complementary subsets of PTCs defined by local sequence context. This allows us to build interpretable models that accurately predict drug-induced readthrough genome-wide. Accurate readthrough quantification and prediction will empower clinical trial design and the development of personalized nonsense suppression therapies.

## Introduction

Premature termination codons (PTCs) are the cause of 10-20% of Mendelian diseases and the most important mechanism of tumor suppressor gene inactivation in cancer ^1, 2^. PTCs cause the production of truncated versions of proteins, which are typically loss-of-function and sometimes gain-of-function or dominant negatives. Many – but not all – PTCs also cause the degradation of g mRNA transcripts by a process called nonsense-mediated mRNA decay (NMD), strongly reducing production of the wild-type (WT) version of a protein ^3, 4^.

A general therapeutic strategy to alleviate the effects of PTCs would be to promote translational readthrough of the stop codon (Fig. 1a). Effective nonsense suppression therapy would increase expression of full length proteins, reduce production of pathological protein fragments, and reduce the harmful effects of NMD ^5^.

**Fig. 1:**
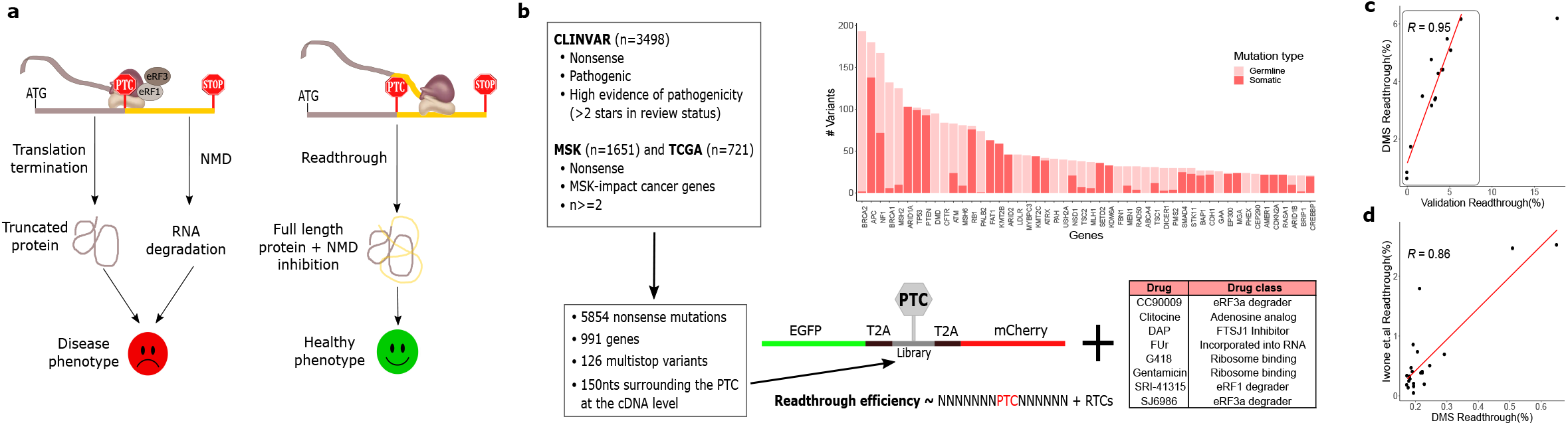
Quantifying readthrough of thousands of pathogenic PTCs. **a**, Pathogenic nonsense mutations relevant to Mendelian diseases and cancer. Readthrough drugs stimulate stop codon recognition by near-cognate tRNAs. **b,** Experimental design, where the most recurrent ∼6000 nonsense mutations in human genetic diseases and cancer where retrieved from ClinVar, TCGA and MSK-IMPACT datasets, cloned in a readthrough reporter, integrated into the genome of HEK293T human cell line and treated with 8 readthrough compounds. A readthrough efficiency value was obtained for each variant-drug pair. Histogram shows the proportion of germline and somatic variants for the top50 represented genes. **c,** DMS measurement correlation with individual measurements in validation experiments, where each variant was individually measured.. **d,** DMS measurement correlation with measurements from a previous study Pranke.et.al [^20^].

Multiple small molecule drugs that promote PTC-readthrough have been discovered, with diverse mechanisms of action promoting the recognition of stop codons by near-cognate tRNAs rather than translation termination factors ^6^. For multiple disease genes, even very moderate readthrough can be sufficient to alleviate disease symptoms in animal models ^7–10^. The extent of readthrough promoted by small molecules varies extensively for different stop codons, with most drugs increasing readthrough of TGA PTCs more effectively than TAG and TAA PTCs ^11–13^. Testing small numbers of mutations has identified sequence features that influence readthrough of particular stops, for example the presence of a cytosine in position +1 after the PTC ^14^ and the presence of an adenine at position –1^15^. To date, the largest survey of drug-induced readthrough tested the compound TLN468 on 40 variants ^16^). Here we generate much richer datasets quantifying nonsense suppression by different drugs. We measure readthrough of ∼5,800 human disease-causing PTCs for eight different readthrough-promoting compounds (henceforth referred to as drugs). We find that the drugs vary substantially in their efficacy and also in the identity of the PTCs that they most effectively promote readthrough of. We identify multiple local sequence determinants that predict PTC readthrough efficacy and show that these determinants differ across drugs. Using these sequence determinants we are able to build models that predict readthrough efficacy by the best performing drugs with very good performance genome-wide. We make these models available as a resource to allow the most effective readthrough-promoting drugs to be identified for all possible PTCs in the human genome. Our data and models show that clinical trials of nonsense suppression therapies have been badly designed, using patient-drug combinations that are unlikely to be effective. Combining data generation at scale and model building to accurately predict drug-induced readthrough will improve the design of clinical trials and facilitate the development of personalized nonsense suppression therapies.

## Results

### Quantifying readthrough of thousands of pathogenic PTCs

To quantify drug-induced readthrough of diverse PTCs we constructed a library containing 3,498 PTCs that cause Mendelian diseases reported in Clinvar ^2^ and 2,372 recurrent somatic PTCs in cancer genes (721 from TCGA ^17^ plus 1,651 from MSK-IMPACT ^18^) (Fig. 1b). We cloned each PTC with 150 nts of surrounding sequence context (75 nt upstream and 75 nt downstream of the PTC) into a dual fluorescent protein reporter, where readthrough causes expression of a downstream mCherry protein. The PTC library was integrated as a single copy per cell into a genomic landing pad locus ^19^ in HEK293T cells. Cells were treated with doxycycline to induce expression of the PTC-containing transcripts, readthrough-inducing drugs were added after 24hrs, and 48hrs later the cells were sorted into 3 to 6 bins according to their mCherry fluorescence. Genomic DNA was then extracted, amplified and sequenced (Extended Data Fig. 1a). The distribution of each PTC across the bins allows us to quantify its readthrough. The resulting efficiencies were highly correlated across replicates and also with individual measurements (up to a measurement ceiling of ∼6%, Fig. 1c). Our measurements also correlate very well with quantifications made using a different technique in a different laboratory ^20^ (Fig. 1d).

### Readthrough efficiency varies extensively across drugs and PTCs

In total, we quantified the readthrough induced by 8 different drugs: CC90009 ^21, 22^, clitocine ^23^, DAP ^12^, gentamicin ^24, 25^, G418 ^24, 25^, SJ6986 ^21^, SRI-41315 ^26^ and 5-Fluorouridine (FUr) ^8^ (Extended Data Fig. 1b). These comprise different classes of small molecules. G418 and gentamicin bind to the decoding center of the small ribosomal subunit; SRI, SJ6986 and CC90009 are eRF1/3 inhibitors; DAP interferes with the activity of a tRNA-specific 2′-O-methyltransferase (FTSJ1); whereas clitocine and FUr are nucleotide analogs that are incorporated into the mRNA. Considering all PTCs in the library, the median readthrough varied across drugs from 0.08% (gentamicin) to 1.32% (SJ6986). However, each drug promoted stronger readthrough of a subset of PTCs, with the median readthrough of the top-10% of variants varying from 0.51% (gentamicin) to 4.28% (DAP). Readthrough distributions were unimodal with a long upper tail for seven drugs, whereas clitocine treatment resulted in a bimodal distribution of readthrough percentages (Fig. 2a). In the absence of drugs, only a very small number of PTCs (n=47) gave quantifiable readthrough (>0.5%).

**Fig. 2:**
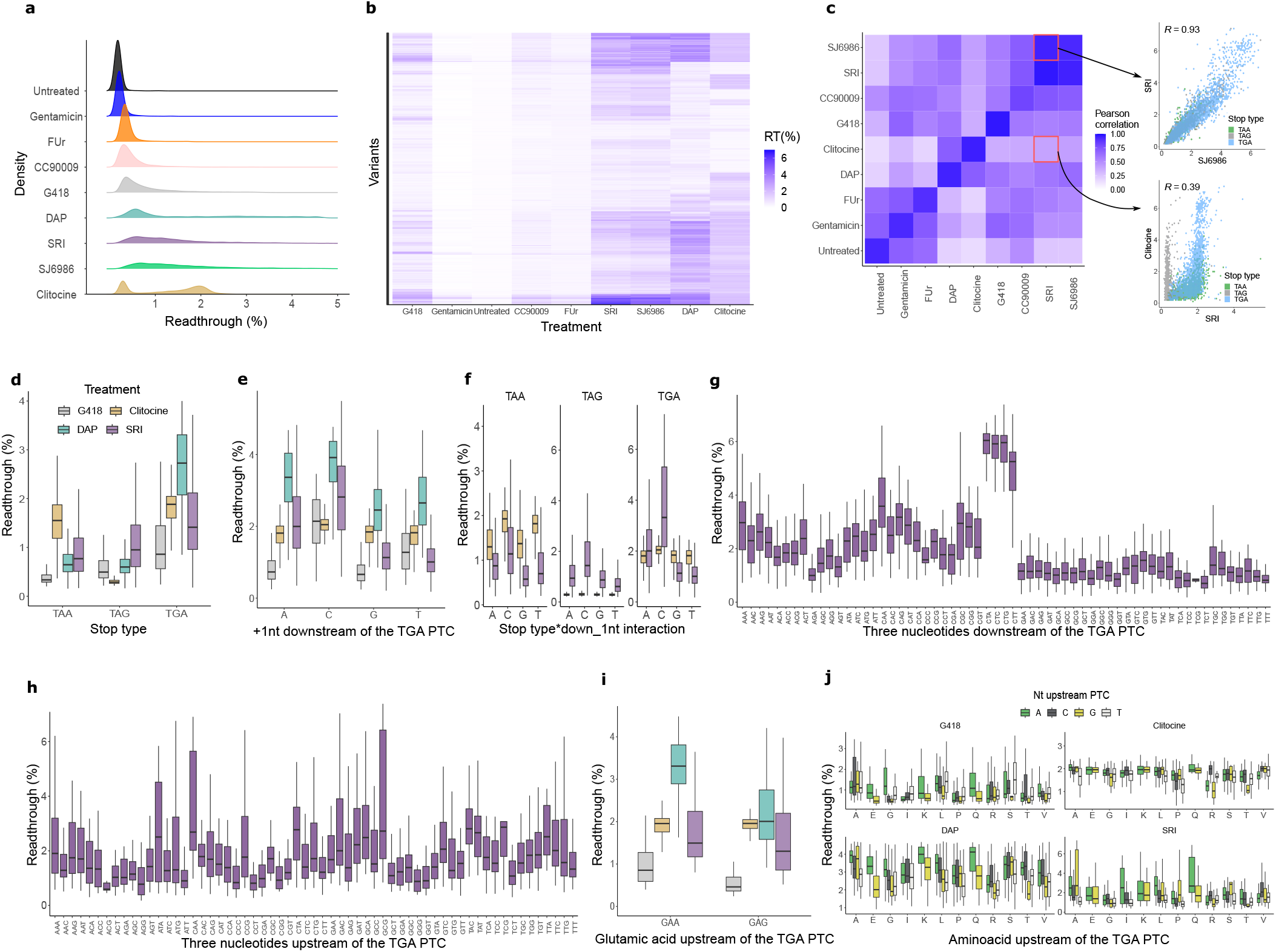
Sequence features explain the readthrough variability across PTCs and drugs. **a**, Readthrough distributions across drugs. **b,** Readthrough efficiency of all variant-drug combinations. **c,** Inter-drugs correlation in RT of variants. Correlation values between the same drug represent the inter-replicate correlation. An example of a high-correlated (SRI and SJ6986) and low-correlated (SRI and clitocine) drug pairs are shown, colored by stop type. **d-i**, Effect of the sequence feature (x-axis) on readthrough efficiency (y-axis), colored by drug. The top and bottom sides of the box are the lower and upper quartiles. The box covers the interquartile interval, where 50% of the data is found. The horizontal line that splits the box in two is the median. Only variants where the stop codon is TGA are shown (except for **d** and **f**, where all stop codon variants are shown). The sequence features are stop codon identity **(d)**, the nucleotide in position +1 downstream of the PTC **(e)**, same as **e** but stratified by stop codon **(f)**, the nucleotides in +1, +2 and +3 positions downstream of the PTC **(g)**, the nucleotides in –1, –2 and –3 positions upstream of the PTC **(h)** and showing variants with a glutamic acid upstream of the PTC clustered by the codon **(i)**. **j,** Effect of aminoacids encoded by A-ending codons on readthrough efficiency across drugs. The nucleotide upstream of the PTC is colored.

The readthrough profiles of the different drugs are, in most cases, only moderately correlated (Fig. 2c, Extended Data Fig. 1e). One exception is SRI and SJ6986, which both inhibit eRF1/3 ^21, 26^ and induce readthrough of a highly correlated set of PTCs (r=0.93, Fig. 2c). The effects of other drugs are much more distinct. Clitocine and SRI, for example, both elicit high readthrough of many PTCs, but their effects are only weakly correlated (r=0.39, in comparison to the inter-replicate correlations of r=0.94 and r=0.96 for the two drugs (Fig. 2c, Extended Data Fig. 1e). Hierarchical clustering of the readthrough profiles of all 5,837 PTCs identifies sets of PTCs with strong readthrough induced by multiple drugs as well as PTCs strongly affected by only one drug (Fig. 2b).

### Stop type and downstream sequence modulate readthrough

To better understand why readthrough of particular PTCs is promoted by particular drugs, we quantified the association between readthrough and 47 sequence features. These included the stop codon type, the adjacent downstream and upstream nucleotides (up to 8 nucleotides away), several codon-related metrics and general features such as GC content and RNA secondary structure propensity (Extended Data Fig. 1f, Extended Data Table.6).

Consistent with previous observations ^27–29^, drug-induced readthrough is much stronger for particular types of stop codon. However, this varies extensively across drugs. For example, for G418 and SRI, the efficiency of readthrough is TGA>TAG>TAA, whereas for clitocine it is TGA=∼TAA>>TAG, and for DAP it is TGA>>TAG=∼TAA (Fig. 2d, Extended Data Fig. 2c). Drugs with the same direction of effect can also have different magnitudes of effects. For instance, both DAP and SRI stimulate TGA>TAG, but the fold-change is different (4.65-fold and 1.64-fold, respectively).

To control for the strong effect of the stop codon types, in the following sections we focus on TGA variants because they trigger highest readthrough across all drugs (conclusions for TAG and TAA are similar and presented in Extended Data Table 2, with differences pointed out in the text). The three nucleotides immediately after a stop codon have been previously reported to modulate readthrough efficiency in the absence of drugs ^27, 30^. Consistent with this, we see a strong effect of the downstream sequence (+1, +2 and +3 nucleotide) on drug-induced readthrough. However, as for stop codon preferences, how the downstream sequence modulates readthrough is drug-specific. Readthrough by all drugs is modulated by the nucleotide immediately after the stop codon (Fig. 2e, Extended Data Fig. 2d). For all drugs, the preferred +1 nucleotide is C. But the rest of the nucleotides show distinct preferences across drugs. For example, T is the most readthrough insensitive +1 nucleotide for SRI and SJ6986, whereas it is the second most sensitive for G418 and gentamicin. Interestingly, drugs with a similar preference for particular stop codons can differ in their preference for the +1 nt. G418 and SRI both promote readthrough of TGA codons over TAG and TAA, but G418 preferences at +1nt are C>T>G=∼A and SRI’s are C>A>G>T. The different drugs also show quantitative differences in their +1nt preferences. For example, readthrough by all drugs is greater for C (n=542) than G (n=1074) at the +1nt but this preference is stronger for SRI, SJ6986 and G418 (3-, 2.5– and 2.8-fold, respectively) than for clitocine and DAP (1.2– and 1.5-fold, respectively) (p<0.01, two-sided Student’s t-test) (Fig. 2e).

Readthrough is also modulated by the +2 and +3 nts, and the effects also differ across drugs (Extended Data Fig. 2a, Fig. 2g). For example, readthrough by SRI is stronger for the downstream nts CAA (n=49) than for CAC (n=28) (1.5-fold, p=4e-05) and for ACT (n=40) than for ACA (n=41) (1.4-fold, p=0.001). In contrast, readthrough of stops with the downstream sequences ACT or ACA is similar for clitocine (p=0.5). CT is the top readthrough-promoting dinucleotide at the +1 and +2 positions for all drugs (CT (n=166) vs all other dinucleotides (n=2558) shows 3.3-fold for SRI, 2.4-fold for SJ6986, 1.4-fold for DAP; all significant at p<=2e-16) except for G418, for which the readthrough is similar across all variants with C in position +1 but CA displays slightly higher readthrough than CT (CT (n=166) vs CA (n=188) shows 1.8-fold for SRI and 1.4-fold for SJ6986, but 0.8-fold, for G418; all significant at p<=1e-08) (Fig. 2g, Extended Data Fig. 2a). Note that the effect of the +3 position depends on the nucleotides at +1 and +2 positions. For instance, an A at +3 position is the top readthrough nucleotide for SRI when positions +1 and +2 are AA (n=84, n=206) (1.2-fold, p=0.005) or CA (n=49, n=139) (1.3-fold, p=0.001), whereas A at +3 is the least readthrough-promoting nucleotide when preceded by AG (n=49, n=138) (0.6-fold, p=3e-09) and AC (n=41, n=80) (0.8-fold, p=0.01).

We identified a stop codon-dependent effect of the downstream nucleotides (Fig. 2f, Extended Data Fig. 2e), indicating genetic interactions between neighboring nucleotides. In the SRI dataset, for TAA stop codons, at position +1 T>G (1.2-fold, p=0.002, n(T)=213, n(G)=407) whereas for TAG stop codons it is the opposite and at +1 G>T (1.3-fold, p=7e-09, n(T)=322, n(G)=515). Also, at position +1 A=∼G for TAG stop codons (1.1-fold, p=0.004, n(A)=477, n(G)=515), but A>G for TGA stop codons (1.8-fold, p<2e-16, n(A)=766, n(G)=1074). C at +1 position has more efficient readthrough than the other 3 nucleotides in all 3 stop codon contexts ^27, 28^.

### Upstream sequence modulates readthrough

Previous studies in bacteria ^31^, yeast ^32, 33^, and mammalian cells ^34, 35^ have shown that the codons upstream of a stop codon can also modulate readthrough under drug-free conditions. Clustering sequences in our library by the upstream codon reveals upstream preferences for each of the drugs (Fig. 2h, Extended Data Fig. 2b). For instance, under SRI treatment the codons encoding the amino acids P, G and I (n=193) display low readthrough, as opposed to Y– and Q-encoding codons (n=459), which drive high readthrough (1.8-fold in Q,Y vs P,G,I, p<2e-16) (Extended Data Fig. 2f).

Similarly to the downstream sequence, these effects are drug-specific (Extended Data Fig. 2b). For instance, under SRI and G418 treatment, both GAA (n=83) and GAG (n=78) codons immediately before the stop display similar readthrough (1.1-fold, p=0.3), whereas when treated with DAP, readthrough of GAA is greater than GAG (1.5-fold, p=2e-12) (Fig. 2i, Extended Data Fig. 2g). Similarly, CGC (n=25) is more sensitive than CGG (n=20) in clitocine (1.7-fold, p=2e-06) but they undergo similar readthrough under G418 treatment (1.2-fold, p=0.2) (Extended Data Fig. 2i,j). These examples show that readthrough differs for different codons encoding the same amino acid. For instance, for SRI readthrough also differs for others codons encoding arginine (CGC>AGG, 2-fold, p=5e-04, n(CGC)=25, n(AGG)=33, Extended Data Fig. 2h), alanine (GCA>GCT, 1.9-fold, p=2e-05, n(GCA)=47, n(GCT)=46), valine (GTC>GTA, 1.5-fold, p=0.001), and leucine (TTA>CTT, 1.7-fold, n(TTA)=30, n(CTT)=77, p=8e-05) (Fig. 2h). To gain more insight into the effect of the upstream RNA sequence, we clustered the codons by the identity of the 3rd nucleotide (Fig. 2j, Extended Data Fig. 2j). Codons ending in A (n=538 vs n=1457) tend to be the top-readthrough promoting codons for all readthrough drugs, except for clitocine, although the effect differs across amino acids (1.1– to 1.3-fold change and p<1e-4 for DAP, G418, SRI, SJ6986 and CC90009).

We found little association between readthrough and GC-content or codon bias indexes (CAI ^36^, tAI ^37, 38^, see Methods and Extended Data Fig. 3a-c), suggesting that the effects of upstream sequence on readthrough are primarily mediated at the mRNA level. Controlling for nucleotide sequence does, however, suggest an additional effect of the encoded amino acid (Extended Data Fig. 2k).

### Readthrough of nonsense mutations in tumor suppressor genes

As evidenced above, there is considerable drug-specific variability in readthrough at the variant level (Fig. 2b). Comparing the top 50 most sensitive variants for each drug shows that each drug maximizes readthrough of a different set of variants (average of 13 variants overlap across all pairwise comparisons, Fig. 3b). Thus the best drug to apply would strongly depend on the particular nonsense mutation causing the disease in each patient.

**Fig. 3:**
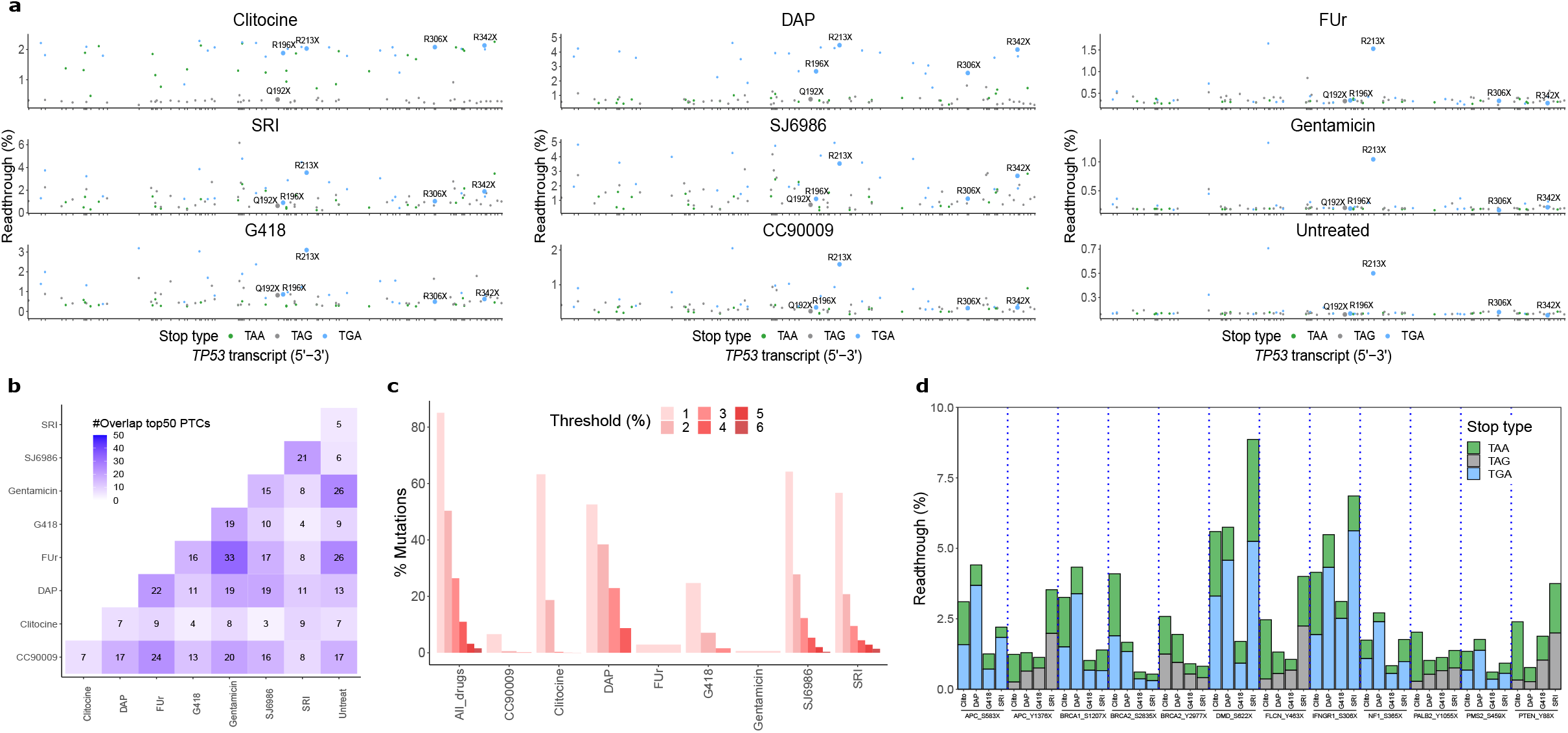
Readthrough sensitive nonsense variants differ across drugs. **a**, Readthrough efficiency, across drugs, for 102 nonsense *TP53* mutations colored by stop codon type. The top-5 most recurrent nonsense mutations in the human tumor genomes are highlighted. **b,** Overlap of the top-50 readthrough sensitive variants for each drug. **c,** Percentage of variants with readthrough over different thresholds for each drug separately and when considering all eight drugs together (“All_drugs”). **d,** Readthrough efficiency for 12 multistop variants across four drugs. Each multistop variant comprises at least two different nonsense mutations (different stop codon identity) observed in the same genomic locus.

As an example of how different drugs promote readthrough of different PTCs we consider the commonly mutated tumor suppressors *TP53* and *PTEN* (Fig. 3a, Extended Data Fig. 3d, Extended Data Table.7). The five most recurrent *TP53* PTCs constitute 44% of all the *TP53* nonsense mutations in the MSK-IMPACT and TCGA datasets (102 different *TP53* nonsense mutations) and are carried by 3% of all cancer patients ^8, 23, 39^. These PTCs show promising readthrough therapy potential. Readthrough of the most prevalent *TP53* nonsense mutation R213X can be substantial: 4.5% with DAP and 3.6% with SRI and SJ6986. Readthrough of the second most frequent PTC, R342X, is strong with clitocine, DAP, SRI and SJ6986 (all >2%). Q192X, the fifth most recurrent mutation, is the only variant in this set insensitive to all treatments (readthrough <1%). In total, readthrough >2% can be achieved by at least one drug for 43/102 PTCs in *TP53*. Considering all 102 PTCs, SJ6986 is the most effective drug for 36 PTCs, DAP for 25 PTCs, clitocine for 21 PTCs, SRI for 14 PTCs and G418 for 6 PTCs in *TP53*.

For *PTEN*, the four most frequent PTCs constitute 36% of PTCs reported in MSK-IMPACT (97 different *PTEN* nonsense mutations were considered here). Of these, the two more prevalent mutations (R130X and R233X) show very similar readthrough profiles with high readthrough with DAP (3.4%, 3.1%) and readthrough also effectively promoted by SJ6986 (2.0%, 2.0%), clitocine (1.7%, 1.8%), and SRI (1.8%, 1.3%) (Extended Data Fig. 3d). In contrast, *PTEN* Q171X and Q245X do not respond to any of the readthrough drugs tested here (readthrough <1%). In total, readthrough >2% can be achieved for 35/97 of all the PTCs reported in *PTEN* in MSK-IMPACT with at least one drug. Considering all 97 PTCs, DAP is the most effective drug for 32 PTCs, clitocine for 31 PTCs, SJ6986 for 22 PTCs and SRI for 12 PTCs.

### The most effective drugs for readthrough of pathogenic variants

The drug-specific readthrough of different variants increases the number of patients potentially treatable by a genetically-informed choice of drug. Considering all 5,837 PTCs in our library, readthrough >2% can be achieved for 50.3% using one of the eight drugs. This is higher than for any individual drug, with >2% readthrough for 38%, 28%, 21%, 19%, 7% and 0.6% of PTCs with DAP, SJ6986, SRI, clitocine, G418 and CC90009, respectively (Fig. 3c). Applying ‘genetically-informed readthrough drug selection’, many mutations display even higher readthrough: RT>3% for 27% of PTCs; RT>4% for 11%; RT>5% for 3.2%, RT>6% for 1.6% (Fig. 3c).

### PTCs in the same position display different readthrough depending on the stop codon type and drug

Our library comprised a total of 240 genomic positions with 2 variants of different stop types (named ‘multistop variants’). The correlation of readthrough between pairs of stop variants ranges from ∼0 to almost 0.85, depending on the drug and stop types being compared (Extended Data Fig. 3e). For instance, in SRI and SJ6986 the TGA variants correlate well in readthrough efficiency with TAA variants but their readthrough is two times higher. With G418, TGA variants are three times more readthrough-sensitive than TAA variants. Other comparisons show very different behavior across stop types (e.g. TAA vs TAG for DAP and clitocine). Examples of different stop type variants in the same genomic position under different treatments are shown in Fig. 3d. For instance, *DMD*_S622X_TGA responds efficiently to DAP but *DMD*_S622X_TAA responds poorly. Other examples include *PTEN*_Y88X in clitocine, *IFNGR1*_S306X in DAP/G418/SRI, and *APC*_S583X in DAP/SRI, amongst others.

### Accurate prediction of readthrough efficiency

Our extensive and quantitative dataset of drug-induced PTC readthrough provides the first opportunity to train and evaluate computational models to predict drug-induced readthrough. We focussed on the 6 drugs that triggered readthrough (>1%) for >3% of PTCs and used logistic regression to train sequence-based genotype-phenotype models (see Methods).

We found that a simple model using four sequence features showed good performance across all 6 drugs (Fig. 4a). The four features included are: a) stop codon type; b) the three nts downstream of the PTC and their interactions; c) the three nts upstream of the PTC and their interactions; and d) the interaction between stop type and the three nts downstream of the PTC. Increasing the downstream sequence context up to +8 nucleotides didn’t improve predictive performance for any of the drugs (Fig. 4c). The correlation between predicted and observed readthrough evaluated by 10 rounds of cross validation (90%-10% training-testing data) was r^2^=0.88 (clitocine), 0.87 (DAP), 0.76 (SRI), 0.76 (G418), 0.71 (SJ6986) and 0.56 (CC90009). Of note, CC90009 is the dataset with highest technical noise (r=0.79 inter-replicate correlation), likely hindering model performance. Training a single model with the data from all six drugs provided poor performance (r^2^=0.39) unless the drug identity was included as a predictive feature with interaction terms between the drug and the additional features (r^2^=0.83 for the pan-drug model) (Fig. 4b). This model explains 94% of the explainable readthrough variance (maximum possible r^2^=0.89 calculated via inter-replicate correlation across all variants and drugs).

**Fig. 4:**
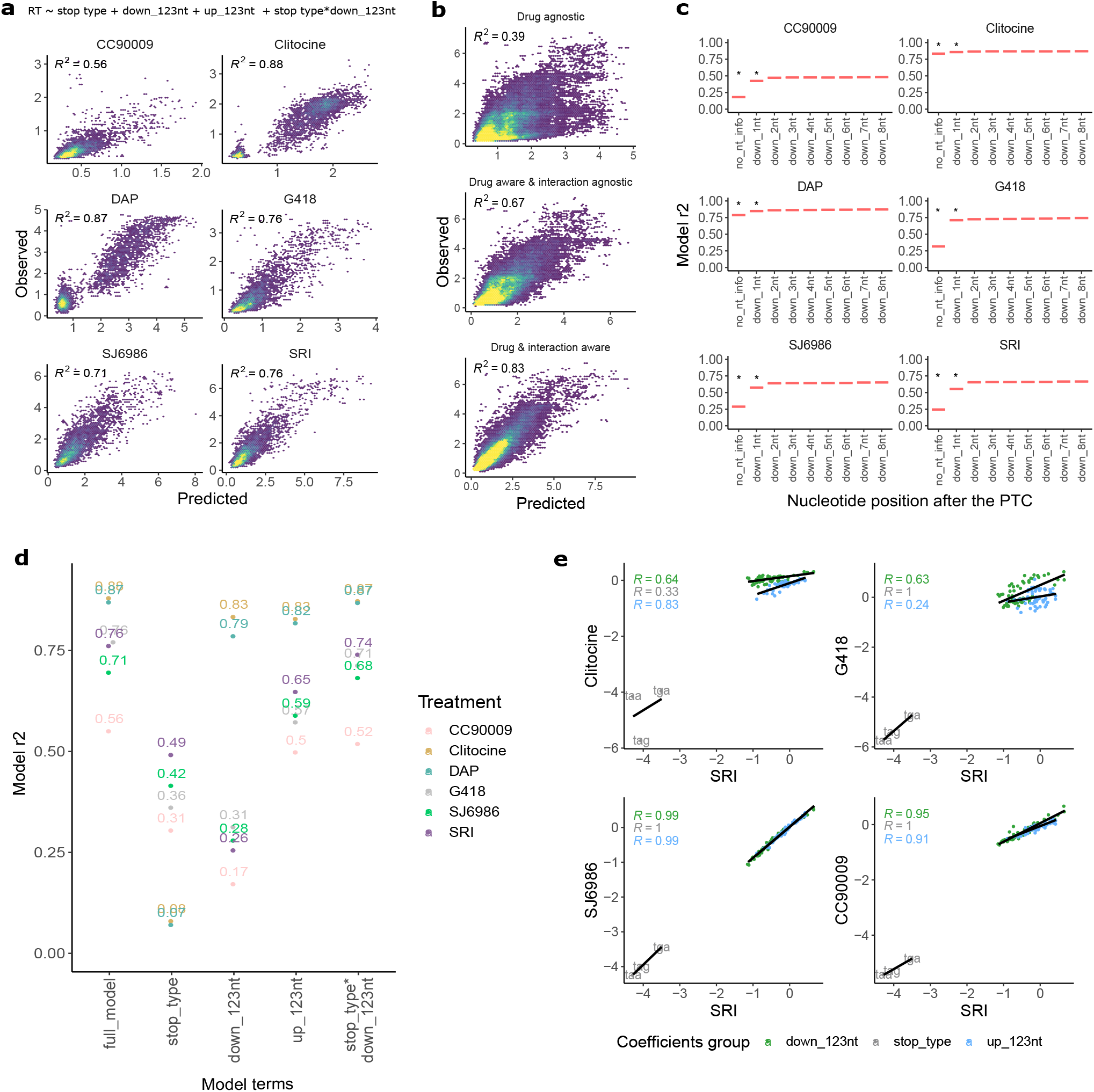
Interpretable models predict readthrough efficiency from sequence context. **a**, Drug-specific models predictive performance in cross-validation. **b,** Pan-drug models performance: drug-agnostic (top), drug-aware but sequence*drug interaction-agnostic (middle), drug and sequence*drug interaction-aware (bottom). **c,** Contribution to model performance of the eight nucleotides downstream of the PTC (by adding one at a time). The fixed predictive variables present in all models are stop codon type and the three nucleotides upstream of the PTC. T-test over 20 cross validation rounds comparing each model (column) to the previous one was used to determine significance (p<0.05). **d,** ANOVA test on the nine drug-specific models. Y-axis shows the drop in model performance upon removal of each predictive variable (X-axis) while retaining all other variables in the model. **e,** Correlation of drug-specific model coefficients (note that for the sake of coefficient interpretability we ran the models without the interaction term stop_type*down_123nts, which incurs only a small decrease of r^2^, ranging between 1-3%). Coefficients are colored by the model feature they belong to: stop codon type, down_123nt and up_123nt. Drugs displaying high correlations respond similarly to the sequence features and consequently, display readthrough in similar subsets of PTCs.

To identify features important for model performance, we removed one variable at a time and calculated the drop in cross-validated r^2^ (Fig. 4d, Extended Data Fig. 4a). Feature contributions quantitatively differ across readthrough drug models. For DAP and clitocine, stop codon type explains most of the variance (80% drop in r^2^ upon stop codon type variable removal) with some contribution of the downstream nts (5-8% drop in r^2^). In contrast, for SRI and SJ6986, the downstream nts are much more important for model performance (50% and 43% drop in r^2^ upon removal, respectively) and the dependence on stop codon type is lower (27% and 28% drop in r^2^ upon removal). For G418 there is a very similar performance reduction upon removing the stop type and downstream nts (40% and 45% drop in r^2^ upon removal, respectively). The upstream nt sequence is less important in general than the downstream nt sequence, but there is still a substantial drop in performance when removing it for SRI, SJ6986 and G418 (11%, 12% and 19% drop in r^2^ upon removal, respectively). The interaction between the stop codon type and the downstream nts is least important for model performance, with r^2^ dropping by 1-3% upon its removal. Analysis of the pan-drug model gives very similar conclusions (Extended Data Fig. 4a).

We also compared the model coefficients for each feature (in a simplified model without the interaction term to aid coefficient interpretability, see Methods) (Fig. 4e). The correlations of the coefficients for each feature between drugs reflects the known mechanisms-of-action. For example, the coefficients for SRI are highly correlated with those for SJ6986 and CC90009 for all three types of features (upstream nts, downstream nts and stop codon), suggesting these drugs show a similar dependence on sequence context. In contrast, although the stop codon coefficients are perfectly correlated between SRI and G418 (r=1), the upstream nt coefficients are only poorly correlated (r=0.24). This indicates that readthrough by SRI and G418 respond very similarly to stop codon type but very differently to changes in the upstream sequence. In contrast, readthrough by SRI and clitocine respond similarly to changes in the surrounding sequence context (r=0.83 and r=0.64 for upstream and downstream nt coefficients, respectively) but very differently to the stop codon type (r=0.33). SRI shows high correlation with SJ6986 and CC90009 across all coefficients, likely due to their shared mechanism of action.

### Improving clinical trial design for nonsense suppression therapy

Our data highlights the highly variable efficacy of readthrough-inducing drugs across different PTCs. However, to our knowledge, only one of 42 phase II-IV clinical trials using readthrough promoting drugs ^40^ used the genetic context of a PTC as an inclusion criteria (Betagov ID: NCT04135495, Extended Data Table 4, Extended Data Fig. 5a). Furthermore, only three of trials made the identity of patient PTCs available in their posterior publications ^41–43^.

We used our readthrough efficiency predictions to evaluate the optimality of the matching between drugs and PTCs in two of these trials using a class of drugs evaluated in our study (Fig. 5a). Trial NCT04140786 used gentamicin and trial NCT04135495 used a gentamicin derivative, ELX-02. However, our data show that effective readthrough of the PTCs present in patients included in these trials is unlikely to have happened. The average readthrough of these PTCs by gentamicin is only 0.2%. In contrast, average readthrough by clitocine and DAP would be 1.8% and 2.9%, respectively, and other PTCs in the same gene would be better choices for a gentamicin trial (Fig. 5a). The most effective drug also varies across patient PTCs in the two other trials for which PTCs were reported (Extended Data Fig. 5b).

**Fig. 5:**
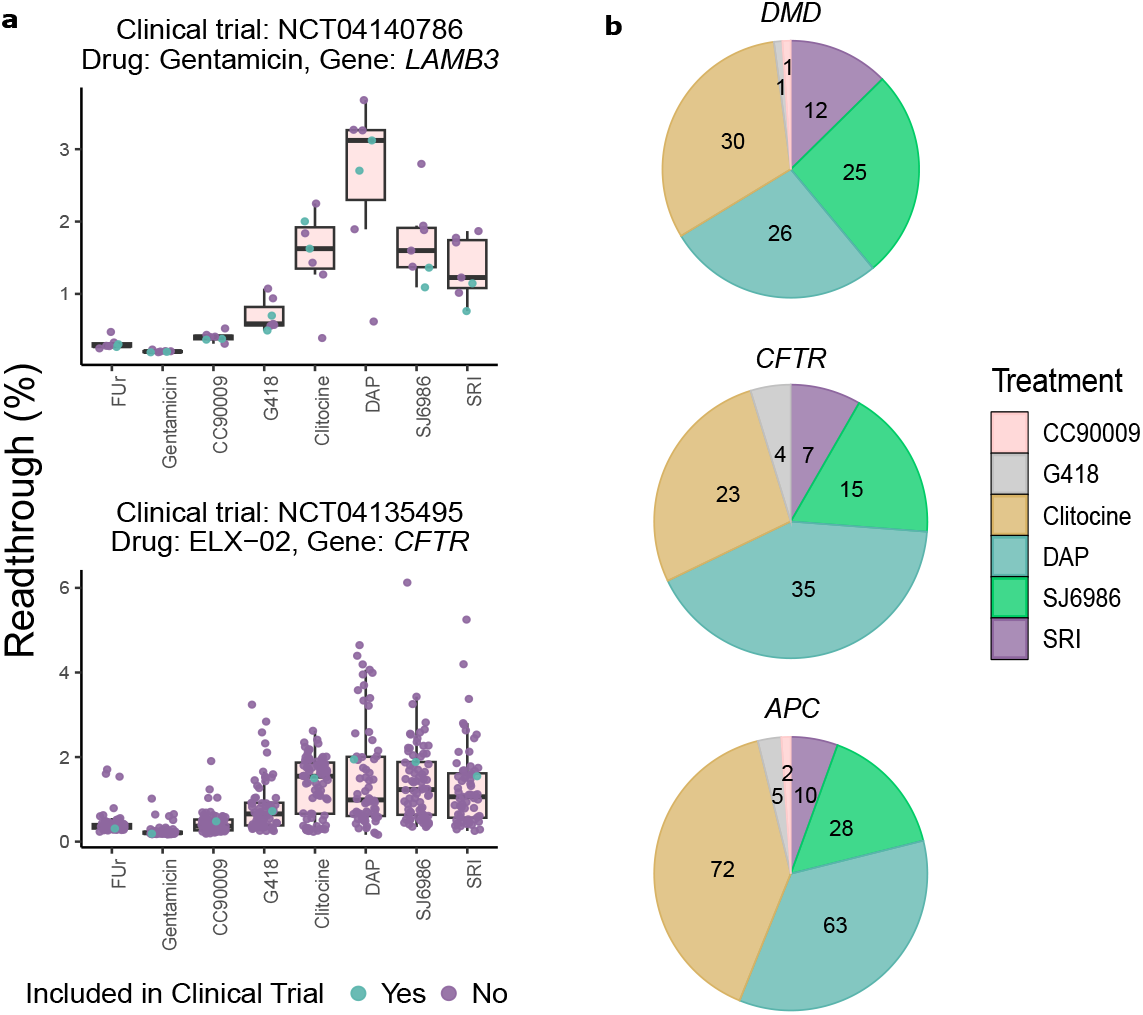
Genetics-informed patient stratification for clinical trials of readthrough drugs. **a**, Our observed readthrough efficiencies of the nonsense variants in two clinical trials where genetic data was supplied (blue), together with the rest of nonsense variants in the same gene (purple). Clinical trial identifier, drug and gene tested are specified in the titles. The top and bottom sides of the box are the lower and upper quartiles. The box covers the interquartile interval, where 50% of the data is found. The horizontal line that splits the box in two is the median. **b,** Number of variants for which each drug displays the highest readthrough efficiency across the top-3 genes commonly tested in the clinical trials, considering all variants in our dataset.

We next considered the three genes most frequently targeted in nonsense suppression clinical trials: DMD, CFTR, and APC. Using our data shows that the most effective readthrough drugs for these genes are clitocine and DAP, but that, in all three cases, a combination of drugs matched to patient PTCs would likely be the most effective trial design (Fig. 5b). For example, of the 95 pathogenic PTCs in DMD, the most effective readthrough is obtained with clitocine for 30 PTCs, DAP for 26, SJ6986 for 25, SRI for 12 and G418 and CC90009 for 1 PTC. None of the best three drugs – clitocine, DAP and SJ6986 – have, to our knowledge, yet been evaluated in clinical trials.

### Nonsense suppression prediction for all PTCs in the human genome

The accurate prediction of drug-induced readthrough by our interpretable model allows us to provide readthrough predictions for every possible PTC in every transcript of the human genome (Fig. 6a). In total, 1-3nt substitutions can introduce 32.7 million stop codons in the 19,061 human protein-coding transcripts (Ensembl v107 genes, hg38 assembly) and we make readthrough predictions for 6 drugs available as a resource named RTDetective that can be visualized along the human genome using the UCSC browser (Figure 6b, Extended Data Fig. 5c-e, doi:10.6084/m9.figshare.23708901).

We estimate that, using these 6 drugs, readthrough of >2% can be achieved for 13M/32.7M (39.6%) of the possible stops in the human genome, with readthrough of >1% possible for 28.6M/32.7M stops (87.3%) (Fig. 6c). The individual drugs are predicted to result in >2% readthrough for 31.4%, 21.3%, 16.2%, 11.7%, 4.3% and 0.02% of PTCs for DAP, SJ6986, SRI, clitocine, G418 and CC90009, respectively. Clitocine is a mid-intensity readthrough drug but spans TAA and TGA stops, inducing 1.5-2% readthrough for many variants but higher readthrough for only a few. Considering all 32.7 million possible PTCs, the most effective drug in 32.8% of cases is DAP, followed by clitocine (30.5%), SJ6986 (29.5%), SRI (6%) and G418 (1.1%) (Fig. 6d).

**Fig. 6:**
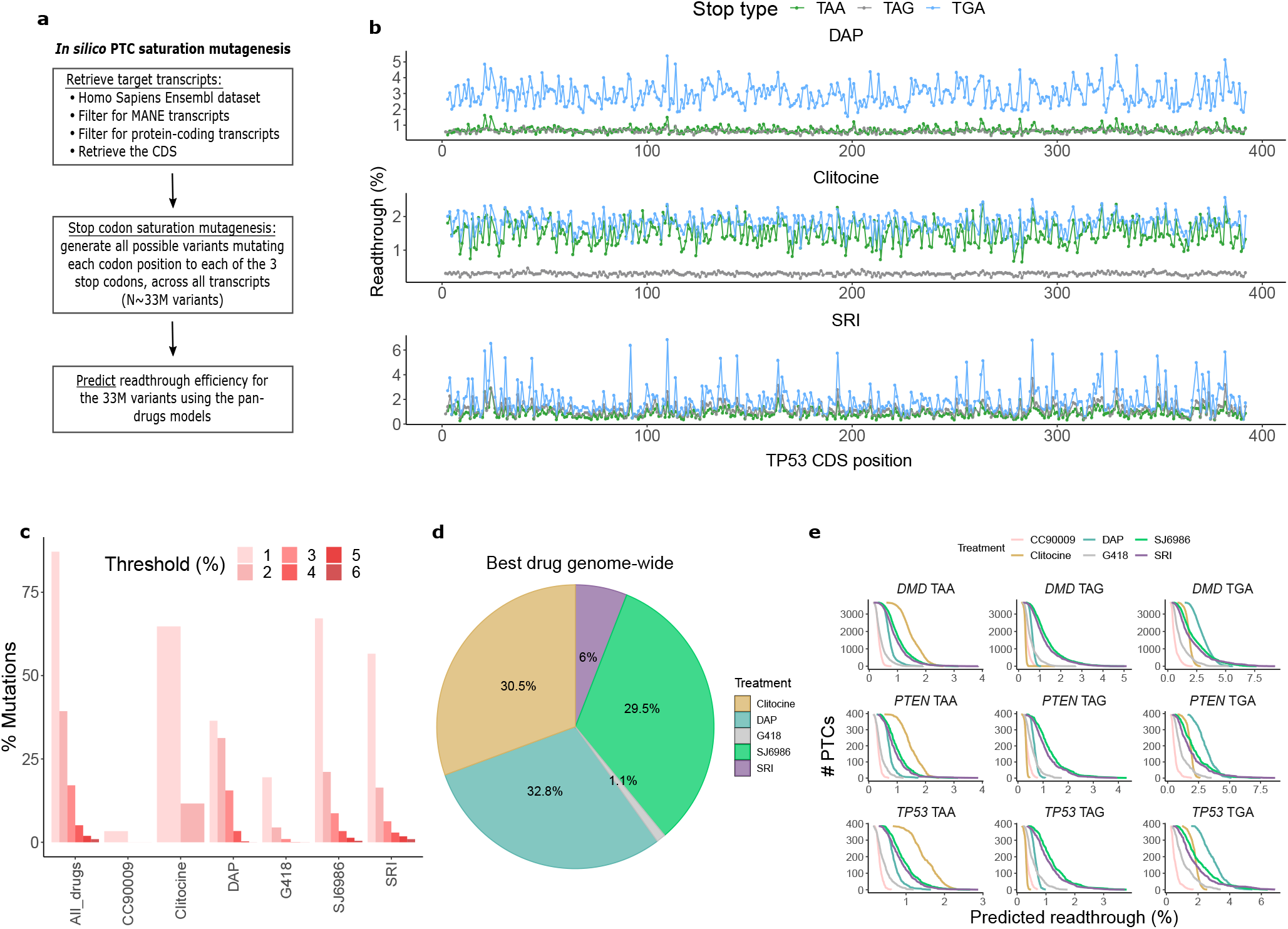
In s*i*lico nonsense saturation mutagenesis of the human genome. **a**, Generation of the comprehensive *in silico* dataset with all possible nonsense mutations. **b,** Readthrough predictions along the coding sequence of *TP53* for each stop codon type. Each row represents a drug-specific readthrough prediction for DAP (top), clitocine (middle) and SRI (bottom). **c,** Percentage of mutations with readthrough over a given threshold (color legend) for each drug. **d,** Number of variants across all possible variants in the human exome for which each drug displays the highest readthrough efficiency. **e,** Cumulative histograms showing number of variants as a function of readthrough efficiency for the genes *DMD* (top), *PTEN* (middle) and *TP53* (bottom), stratified by stop codon type as TAA (left), TAG (center) and TGA (right).

Some drug-stop codon type combinations show promising potential, as illustrated for *DMD, PTEN and TP53* (Fig. 6e). DAP stimulates readthrough >2% for almost 100% of the TGA variants across the three genes. For TAA mutations, clitocine emerges as the best candidate (readthrough>1.5% for ∼50% of PTCs). For TAG mutants, the recently reported eRF3/1 inhibitors SRI-41315 and SJ6986 ^21, 26^ show promise (across DMD, PTEN and TP53, RT >1.5% for 33% and 25% of PTCs for SJ6986 and SRI, respectively; whereas RT >1.5% for <0.1% for Clitocine and DAP). Thus, even for TAG variants that have been considered particularly difficult to suppress, drug-induced readthrough provides a promising therapeutic strategy, provided that the correct drugs are matched to each PTC.

## Discussion

We have presented here the first systematic quantification of drug-induced stop codon suppression comprising >50,000 readthrough measurements in human cells made using 8 drugs. Our massively parallel assay represents a substantial scaling-up of data production compared to previous studies ^14, 16^, generating datasets of sufficient size to train the first models to accurately predict drug-induced nonsense suppression genome-wide.

Our results show that each drug only induces readthrough of a subset of pathogenic PTCs, and our models describe and predict the readthrough specificity of each drug. Whereas for some drugs, such as clitocine and DAP, the type of stop codon is the most important determinant of readthrough, for other drugs, including SJ6986 and SRI, it is the surrounding sequence context. For all drugs, however, accurate prediction requires the stop type and both the upstream and downstream sequence context to be considered.

This diversity of drug responses means that for any particular disease gene – as we have illustrated for *TP53*, *PTEN*, *DMD*, *CFTR* and *APC* – there is no single drug that triggers strong readthrough of all pathogenic PTCs. Rather, effective clinical nonsense suppression will require a panel of drugs, with the appropriate drug selected for each patient according to the identity of the PTC that they carry.

The models that we have trained are deliberately interpretable and relatively simple and yet they explain 94% of variance in readthrough across PTCs. It is possible that black box machine learning models may further improve predictive performance, but model interpretability both aids mechanistic understanding and is highly desirable for models to be used in clinical decision making ^44^.

We envisage that accurate genome-scale prediction of drug-induced readthrough will greatly improve clinical trial design and the development of personalized nonsense suppression therapies. To date, trial designs have nearly all ignored the large variation in readthrough efficacy across PTCs ^40^, and our data show that this has resulted in suboptimal matching between patient PTCs and drugs. The use of the right drugs but in the wrong patients is, in retrospect, likely to have been an important cause of trial failure.

Moving forward, the power of clinical trials will be substantially boosted by using our approach to stratify patients by their PTCs so that each drug is only tested in the subset of patients in which it will promote strong readthrough and not in patients in which there will be zero or very little readthrough. The data, models and genome-wide predictions we provide can be used for this purpose and our general approach can be used to rapidly quantify the specificities of novel nonsense suppression therapeutics, allowing their clinical efficacy to then be tested in the subset of patients in which they are likely to be most effective.

Taken together, our results show that the specificities of nonsense suppression therapies differ extensively and these specificities can be rapidly learned using high-throughput experiments to allow accurate prediction of drug responses. Looking forward, we believe that the goal should be to develop an expanding portfolio of readthrough drugs with defined and complementary specificities such that effective and specific nonsense suppression therapy can be achieved for any pathogenic stop codon in the human genome.

## Methods

### Library design

Genetic disease and cancer germline variants were retrieved (n=3498) from ClinVar database, where all pathogenic nonsense variants whose review status was two or more stars were included (a final filtering step was carried out to decrease the penetrance of overrepresented genes such as BRCA2). Somatic cancer variants were obtained from MSK-Impact and TCGA databases with at least two entries in either dataset (n=2372). Also, 15 genomic sequences of viruses with reported readthrough events were included in the dataset. Finally, a control WT non-stop TP53 variant was included to use as a 100% readthrough control, in order to normalize the expression of the PTC variants and get estimates for the percentage of WT protein expression for each variant. The 150nt spliced-mRNA context of each nonsense variant (MANE transcript isoform) was retrieved from Ensembl (v104) in order to preserve a large sequence context (75nts upstream and 75nts downstream).

### Readthrough reporter

We designed a double fluorescent reporter plasmid (pIT092) to quantify readthrough (sequence available in Supplementary File 1). The plasmid encodes a single transcript that contains the ORFs of EGFP, T2A (x2) and mCherry from 5’-3’, respectively. The library oligopool was cloned in frame between the two T2A sequences. T2As allow the independent folding of the fluorescent proteins and prevent undesired effects of the variable sequence on their folding and stability. In a normal translation event termination occurs in the PTC of the library, protecting mCherry from translation. However, if readthrough occurs, the ribosome extends elongation until the mCherry stop codon, translating the mCherry protein along the way. Hence, mCherry fluorescence is proportional to readthrough efficiency, and we used it as our assay readout. EGFP is used to filter out those cells that either don’t have EGFP or have unexpectedly high levels of EGFP. These cells are likely to have either aberrant cloning, out-of-frame integration, promoter mutations, promoter silencing, transcript-stability mutations, etc, and might be misleading if included in the assay. For some of the treatments, we detect a slight EGFP increase in the mCherry+ population, suggesting a readthrough-mediated transcript stabilization either via NMD inhibition or translation-mediated mRNA protection. In those cases, the mCherry increase (in mCherry+ vs mCherry-populations) is higher than the EGFP increase, proving that the drugs are increasing mCherry signal via readthrough. pIT092 is suitable for genomic integration using the HEK293T landing pad system (Matreyek et al., 2020), which ensures that each cell integrates only one variant providing a direct genotype-phenotype linkage. The vector contains BxBI-compatible attB sites that allow recombination into the genomic landing pad of HEK293T landing pad cells. After genomic integration into the landing pad locus, the ORF sequence is placed right downstream of a tetracycline induction cassette, allowing its expression when doxycycline is added to the media.

### Library cloning

Oligos were ordered as an oligopool to Twist Biosciences containing the variable part (library) and two constant sequences in both for the PCR amplification. The oligopool was PCR-amplified for 14 cycles using primers oIT204 and oIT340 (Extended Data Table.5). The oligo pool was cloned between the EGFP-T2A and T2A-mCherry open reading frames (ORFs) of pIT092 using Gibson Assembly. The library was electrotransformed using Neb10 Electrocompetent bacteria and grown in 100mL overnight culture. Library complexity and representativity of the variants was estimated by plating a small amount of the transformation reaction and extrapolating the total number of transformants. Individual clones were Sanger sequenced to confirm the expected structure and diversity.

### Stable cell line generation

To generate the cell line, we used the HEK_293T landing pad cell line generated by Matreyek et al., 2020 (TetBxB1BFP-iCasp-Blast Clone 12 HEK293T cells), which allows the stable single-copy integration of variants in the genome. Mutational libraries cloned into the landing pad compatible construct (pIT092) are co-transfected (1:1) with a BxBI expression construct (pCAG-NLS-Bxb1) into the HEK293T landing pad cell line using lipofectamine3000 according to the manufacturer’s instructions in three T150cm2 flasks. This cell line has a genetically integrated tetracycline induction cassette, followed by a BxBI recombination site, and split rapalog-inducible dimerizable Casp-9. Cells were maintained o in DMEM supplemented with 10% FBS tetracycline-free without antibiotics. 2 days after transfection, doxycycline (2 μg/ml, Sigma-Aldrich) was added to induce expression of the library (recombined cells) or the iCasp-9 protein (no recombination). 24h later, 10nM rimiducid (AP1903, Selleckchem) was added to the cells. Successful recombination frameshifts the iCasp-9 out of frame. However, non-recombined cells express iCasp-9 which dimerizes in the presence of Rimiducid and induces apoptosis. One day after rimiducid treatment, the media was changed back to DMEM + Doxycycline, and cells were maintained in culture for the following 5 days to obtain a large volume of cells for downstream experiments and cryostorage.

### Readthrough compounds

We tested a panel of twenty compounds reported to have readthrough activity (Extended Data Table.1), in our library-integrated HEK293T landing pad cells. If a drug induces readthrough, the FACS profile would be different than the vehicle-treated cells, specifically, we would observe an increase in the mCherry+ population (Extended Data Fig. 1b). All drugs were tested at six or more different concentrations ranging along orders of magnitude, to ensure that a negative result was not due to a concentration related problem. 4×10e5 library-integrated HEK293T landing pad cells were seeded in 6-well plates and treated with doxy to induce the expression of the transcript, and after 24hrs the drug was added to the medium. Readthrough was measured 48h after treatment with BD LSRFortessa™ Cell Analyzer as described above. Eight drugs, named SRI, Clitocine, SJ6986, DAP, G418, Gentamicin, CC90009 and FUr, were validated whereas the remaining twelve were not able to trigger detectable readthrough in our system at any of the tested concentrations. We did toxicity titrations for the eight positive drugs (Extended Data Fig. 1c). 2×10e4 cells were plated in 96-well plates and CellTiter-Glo® Luminescent Cell Viability Assay (Promega) was used to quantify cell viability 48h after drug or vehicle treatment using a Tecan Infinite M Plex plate reader (Tecan, Switzerland). For each drug, the concentration that didn’t decrease cell viability more than 25% and exhibited the highest readthrough was selected.

The validated drugs comprise different classes of small molecules. G418 and Gentamicin bind to decoding center of the small ribosomal subunit; SRI, SJ6986 and CC90009 are eRF1/3 inhibitors; DAP interferes with the activity of a tRNA-specific 2′-O-methyltransferase (FTSJ1); whereas Clitocine and FUr are nucleotide analogs that get incorporated into the mRNA. Some were reported as drugs decades ago and their readthrough potential is supported by extensive literature (Gentamicin, G418) and tested in several clinical trials (most of them with disappointing and confusing outcomes). In contrast, others have been recently described as drugs and little is known about their readthrough stimulatory potential.

### Fluorescence activated cell sorting (FACS)

Cells were grown on standard culture plates in DMEM supplemented with 10% FBS tetracycline-free and without antibiotics. They were split before reaching confluency to maintain cell health. Cells were detached with Trypsin, spun down and washed with PBS. For the sort-seq experiments, cells were treated with doxy to induce the expression of the transcript, and after 24hrs the drug was added to the medium for 48h more. We used high volumes of cells to ensure that each variant was represented >100 times in the cell population. Before sorting, the same amount of cells as those sorted in each bin were withdrawn for sequencing as a pre-sorting control.

Cells were sorted on a BD Influx (TM) Cell Sorter. Cells were gated by forward scattering area and by side scattering area to retain whole cells, forward scattering width and height to separate discard aggregates, and by DAPI-staining to retain only recombined and alive cells. EGFP and mCherry fluorescence were excited with a 488nm and 561nm lasers and recorded with a 530/40 BP and 593/40 BP channels, respectively. EGFP+ cells were sorted based on mCherry expression into four populations. For most of the populations, 400K cells were sorted. However, for some minor populations representing <2% of the total population, we sorted less cells (100-200K). The percentage of cells in each bin population was used for normalization during sequencing analysis (see below). Two replicates were performed for each experiment.

### DNA extraction

Genomic DNA extraction was carried out following the DNeasy Blood&Tissue Kit (Qiagen), and resuspended in 80 ul of MilliQ.

### Sequencing library preparation

The sequencing libraries were constructed in three consecutive PCR reactions. The first PCR intends to amplify the library fragment from the genomic DNA pool, without amplifying the remaining plasmid from transfection. It uses a forward (oIT314) primer annealing in the landing pad outside of the recombined sequence, and reverse (oIT205) primer annealing at 3’ end of the library fragment. This ensures that plasmid DNA is not amplified since it lacks the annealing site for the Fwd primer.The second PCR (PCR2) was designed to insert part of the illumina adapters and to increase the nucleotide complexity of the first sequenced bases by introducing frame-shift bases between the adapters and the sequencing region of interest. The third PCR (PCR3) was necessary to add the remainder of the Illumina adapter and demultiplexing indexes. All PCRs were run using Q5 Hot Start High-Fidelity DNA Polymerase (New England Biolabs) according to the manufacturer’s protocol.

All genomic DNA extracted from each bin was used as template for PCR1, and amplified using 25 pmol of primers oIT205 and oIT314. Annealing temperature was set to 66°C, extension time to 1 minute and number of cycles to 25. Because high volumes of genomic DNA inhibit PCR reaction, we aliquoted each sample in 8 PCRs and run them in 96 well plates. Excess primers were removed by adding 0.04 uL of ExoSAP-IT (Affymetrix) per uL of PCR1 reaction and incubated for 20 min at 37°C followed by an inactivation for 15 min at 80 °C. Then, PCRs of each sample were pooled together and purified using the MinElute PCR Purification Kit (QIAGEN) according to the manufacturer’s protocol. DNA was eluted in MilliQ to a volume of 20ul.

2ul of PCR1 product were used as template for PCR2, together with 25 pmol of pooled frame-shift primers (oIT_ILL_204_mix and oIT_ILL_205_mix) (Extended Data Table.5). The PCR reactions were set to 66°C annealing temperature, 15 seconds of extension time and run for 8 cycles. Excess primers were removed by adding 0.04 uL of ExoSAP-IT (Affymetrix) per uL of PCR1 reaction and incubated for 20 min at 37 °C followed by an inactivation for 15 min at 80 °C. The PCRs of each sample were purified using the MinElute PCR Purification Kit (QIAGEN) according to the manufacturer’s protocol. DNA was eluted in MilliQ to a volume of 10ul.

2ul of PCR2 products were used as template for PCR3. In this second PCR the remaining parts of the Illumina adapters were added to the library amplicon. The forward primer (oIT_GJJ_1J) was the same for all samples, while the reverse primer (oIT_GJJ_2J) differed by the barcode index, to be subsequently pooled together and demultiplex them after deep sequencing (Extended Data Table.5). 8 cycles of PCR2s were run at 62°C of annealing temperature and 25 seconds of extension time. All reactions from the same sample were pooled together and an aliquot was run on a 2% agarose gel to be quantified. After quantification, samples with different Illumina indexes that were sequenced together in the same flow-cell were pooled in an equimolar ratio, run on a gel and purified using the QIAEX II Gel Extraction Kit. The purified amplicon library pools were subjected to Illumina 150bp paired– end NextSeq500 sequencing at the CRG Genomics Core Facility.

### Sequencing data processing

FastQ files from paired-end sequencing of all experiments were processed with DiMSum (reference https://github.com/lehner-lab/DiMSum) to obtain the read counts for each variant. DimSum applies stringent quality filters to discard low quality reads, reads with sequencing errors, etc. to ensure that only high-quality reads are used for downstream analysis.

The DimSum output read count tables were used to calculate readthrough estimates for each variant as follows:

The read count table provides the distribution of each variant among the different sorting gates. Since gates harbored different percentages of the general population, they had to be sorted for different times in order to get the same number of cells in each bin, forcing us to calculate the mCherry distributions in relative numbers. The distribution of each variant is generated by calculating its proportion of reads in each sorting gate (j). All reads of a given variant in a given sorting gate (rj) are a) divided by the total number of reads of that gate (Rj) yielding a normalized reads value b)assigned a first fixed value (*pc*) corresponding to the % of cells of the total population that belong to that gate and c) a second fixed value (*fv*) corresponding to the mean mCherry signal of the gate, and then it’s averaged by the total number of normalized reads across gates (N). In steps a) and b) we are simply calculating the percentage of reads of the total population belonging to each variant in each gate 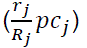. The resulting averaged value corresponds to a normalized mean mCherry value for each variant. Then, by performing the ratio between the mCherry value of each variant and the mCherry value of the WT non-stop variant, the readthrough percentage for each variant (RTp, % of WT protein expression) is calculated. Standard deviation from the two replicates is used as the error measure. RTp values for all variants and drugs are included in Extended Data Table.2.

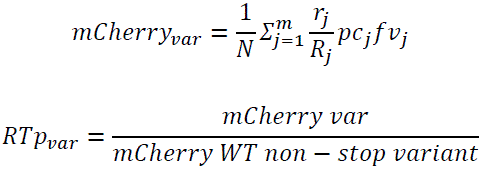

### Upper bound on sort-seq readthrough estimations

Validating the library readthrough values for 15 variants using single variant genome integration by flow cytometry (See <u>single</u> <u>variants validation</u> paragraph below) allowed us to assess the performance of our sort-seq readthrough assay (Fig. 1c). Correlation value (r=0.95, Pearson) is very high when removing the outlier (correlation on 14/15 variants). The inclusion of this outlier drops the correlation to 0.8.

The disagreement in readthrough values between sort-seq and validation in this variant responds to a well-known sort-seq related problem^45^. Variants with readthrough distributions smaller than the sorting gate width are more likely to be miscalled. Our gates were designed in the logarithmic scale, meaning that high-readthrough gates have larger widths than low-readthrough gates. This holds especially true for the highest gate, which was designed to include all cells above the top-edge of the second-highest gate, in order to not lose any high-readthrough cells. In turn, only very high-readthrough variants suffer from the problem described above and this results in our assay having a 6% readthrough upper bound, preventing us from quantifying the exact readthrough efficiency of variants with readthrough>6% (Fig. 1c). How many variants in our assay display readthrough>6%? We computed the percentage of variants in each treatment that had more than 90% of reads in the highest gate. In most of the treatments, this number is around 1% (1.54, 1.31. 0.22, 0.57, 0.41, 1.04, 0.54 and 1.73; for DAP, SRI, Clitocine, SJ6986, CC90009, G418, Gentamicin and FUr, respectively).

### Single variant validation experiments

To validate the assay, we set out to individually measure the readthrough of 15 variants. We selected 15 variants spanning the whole dynamic range when treated with SRI plus the non-stop WT control, and individually cloned and integrated them into HEK_293T_LP, yielding 16 stable cell lines each expressing a different variant. Cells were treated with SRI at the same concentration than used in the DMS assay (7.5uM), and EGFP and mCherry fluorescence were quantified using 530/40 BP and 593/40 BP channels in BD LSRFortessa™ Cell Analyzer. EGFP+ cells were used to calculate the readthrough by multiplying the mean mCherry signal by the percentage of mCherry+ cells, and finally normalizing by the expression of the non-stop WT variant. We termed these readthrough values as RTp_individual (RTp_ind) since they refer to individual measurements of each variant. RTp_individual were correlated against the DMS RTp estimates of the 15 variants to calculate the correlation coefficient between our DMS assay and individually measured readthrough (Extended Data Table.3).

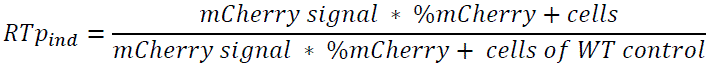

### Sequence features

We listed a large set of sequence features to test their contribution to readthrough variability. Features were chosen based on literature and preliminary results, but we also listed several features for which we had no evidence of their participation in readthrough. All features tested in the predictive models can be found in Extended Data Table.6.

### Model design

We computed the Pearson-correlations (continuous features) or chi-squared (Kruskal-Wallis) values (discrete features) between the listed features and readthrough in our dataset (Extended Data Fig. 1f). We generated models by manually including and excluding the top-correlated features. Along this process, we realized that predicted and observed readthrough were non-linearly related. This non linearity issue was fixed when using general linear models with logit as the link function. In this scenario, we devised a new term referred to as ‘readthrough potential’. Mathematically, it corresponds to the exponential term in the denominator of the model (*Formula 1*). Conceptually, it can be understood as an energetic readthrough value and features additively combine to determine it. The readthrough potential is non-linearly (sigmoid) correlated with the observed/phenotypic readthrough. This is captured in the model by the addition of a sigmoid as a link function between readthrough and readthrough potential.

Models were trained using 90% of the dataset and tested in the remaining 10% of variants (held-out dataset). This process was repeated 10 times, and the mean r^2^ across these cross validation rounds was used as the model performance metric. We sought for those features combinations that maximized model performance (r^2^) without overfitting it (tested by cross-validation).

Our best performing model (Fig. 4a) used the stop type, the three nucleotides downstream of the TGA (down_123nt), the three nucleotides upstream of the TGA (up_123nt) and the interaction between down_123nt and stop_type as predictor variables. This was the top-performing model across all drugs:

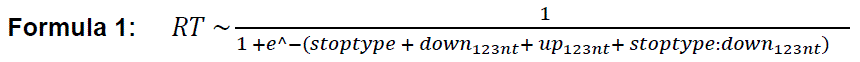

To compare this model to alternative approaches we conducted four analyses::

– Manually test if the inclusion of the stop_type*up_123nt interaction term improved model performance. It did not improve the r^2^ in contrast to the inclusion of the stop_type*down_123nt, which increased r^2^s by 1-3% over the model without interaction terms (Extended Data Fig. 4c).
– Run ElasticNet regularized regression with all features in Fig. 2d allowing for second order interactions. Model performance was not improved for clitocine and slightly improved for the rest of the drugs (r^2^ increased by 1% in CC90009, 1% in DAP, 2% in G418, 3% in SRI and 4% in SJ6986) (Extended Data Fig. 4b).
– We tested whether a simplified encoding of the +/-3nts sequence maintained the high model performance (Extended Data Fig. 4d). We broke down the up_123nts and down_123nts into three terms each: up1, up2, up3 and down1, down2, down3 terms; reducing complexity of the model by cutting down the number of levels from 64 to 12. The other model terms were not modified, and the interaction between stop type and each of the down1, down2, down3 was included. The simplified models consistently show a decreased performance across drugs, suggesting that the interactions between the nucleotides positions are significantly contributing to readthrough and validating our initial model as the best model.
– To ensure that we were not missing effects of nucleotide positions further downstream of the stop, we designed models including each of the eight nucleotide positions downstream of the PTC independently, and checked the contribution of each term in the model as increase in r^2^ (Fig. 4c) by 10 rounds of cross validation, as follows:

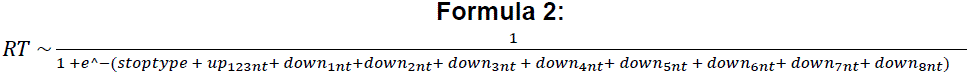 Across drugs the +1 and +2 nts are the only positions contributing to readthrough variability across the dataset. Note that the +3 position is not significant in these models, but it slightly but consistently increases model performance when allowing the interactions with the +1 and +2 positions, suggesting that the effect of +3 position is highly dependent on the nucleotide in +1 and +2 positions (down_123nt consistently improved model r^2^ 1-2% over down_12nts across drugs, data not shown). Importantly, it’s likely that further positions also contribute to readthrough but due to our limited library size and design we can’t neither accurately nor significantly capture the small effects of such positions.
– Lastly, we also tested the effects of the +4 nucleotides downstream considering all possible interactions. We swapped the down_123nt term for the down_1234nt. r^2^ after 10 cross validation rounds was not improved for any of the drugs (data not shown).

#### Models for gentamicin, FUr and untreated conditions

Models trained to predict readthrough for gentamicin (37 PTCs with >1% readthrough) and FUr (153 PTCs with >1%) showed moderate performance (r^2^=0.37 and 0.38, respectively) and models trained to predict readthrough without any drug (17 PTCs >1%) were not predictive (r^2^=0.02). Interestingly, the untreated dataset shows high correlation with both gentamicin and FUr datasets (Fig. 2c, Extended Data Fig. 1e). Thus, gentamicin and FUr, which have a low readthrough-inducing capacity, mostly increase readthrough of those variants which already undergo some basal readthrough (Extended Data Fig. 1e). Downsampling the number of positive examples for the other drugs suggests that this moderate and poor performance likely reflects the small number of positive examples and that improved performance is likely to be obtained by testing a larger set of PTCs (data not shown).

#### Simplified version of the models without the stop_type*down_123nts interaction term

For the sake of coefficient interpretability we also ran the models without the interaction term stop_type*down_123nts, which incurs only a small decrease of r^2^, ranging between 1-3% (Fig. 4d). Cross-validation was used to obtain the mean, standard deviation and significance of the coefficient estimates (Extended Data Fig. 4e-h). Note that coefficients are not in the readthrough percentage scale but in the logit space. Importantly, coefficient sizes must be compared across drugs but within each feature.

#### Pan drug model

We generated a pan drug model by including new terms that would capture the treatment effects on readthrough (combining the data for the six drugs that gave readthrough (>1%) for >3% PTCs, namely clitocine, CC90009, DAP, G418, SJ6986 and SRI). First, a *drug* term was added to the model, which scales the readthrough for each treatment. However, sequence features have different effects depending on the treatment (Fig. 2d-f,i,j). To incorporate such effects into the model, we included 4 interaction terms: *drug:stop_type*, *drug:nt_123*, *drug:up_codon* and *drug:nt_123:stop_type* (in the formula below, the *stoptype* term has been abbreviated to *stop)*. Same as before, model performance was assessed for 10 cross validation rounds.

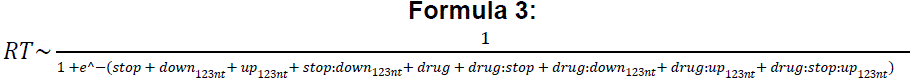

### tRNA Adaptation Index (tAI)

The tRNA Adaptation Index (tAI) is a measure of translational efficiency which takes into account the intracellular concentration of tRNA molecules and the efficiencies of each codon–anticodon pairing^37, 38^. The pairing affinity of each codon-anticodon is specific to each species. The human specific tAI indexes were downloaded from the STADIUM database as of January 2023^46^. tAI for a given sequence was calculated as the mean tAI across all codons of the sequence.

### Codon Adaptation Index (CAI)

The CAI is an estimate of translational efficiency based on the similarity of codon usage of one sequence with regard to the genome codon usage^36^. Codon usage is a metric that computes the genomic abundance of a certain codon compared to the most abundant synonymous codon. The human codon usage table was downloaded from the Codon/Codon Pair Usage Tables (CoCoPUTs) project release as of January 2023^47^. CAI for a given sequence was calculated as the mean CAI across all codons of the sequence.

### *In silico* saturation mutagenesis

We used the general drug model to perform an *in silico* prediction of the readthrough efficiency of all possible nonsense mutations in the human exome, resulting in 32.7 × 10^6^ predictions for the 19,061 protein-coding transcripts (Ensembl v107 genes, hg38 assembly) for each drug. For each codon position of each protein coding transcript, a readthrough efficiency value was estimated for each drug.

## Data availability

All DNA sequencing data have been deposited in the Sequence Read Archive (SRA) with accession number PRJNA996618: http://www.ncbi.nlm.nih.gov/bioproject/996618. The readthrough efficiency predictions have been made available through the Figshare repository at https://figshare.com/articles/dataset/Readthrough_predictions/23708901 and via a digital object identifier (doi:10.6084/m9.figshare.23708901). All readthrough measurements are provided in Extended Data Table 2.

## Code availability

Source code used to perform all analyses and to reproduce all figures in this work is available at https://github.com/lehner-lab/Stop_codon_readthrough.

## Author contributions

I.T., F.S and B.L. conceived the project and designed the experiments; I.T. performed the experiments and analyzed the data; I.T, F.S. and B.L. wrote the manuscript.

## Acknowledgements

Work in the Supek lab is supported by an ERC StG “HYPER-INSIGHT” (757700), Horizon2020 project “DECIDER” (965193), Spanish government project “REPAIRSCAPE”, CaixaResearch project “POTENT-IMMUNO” (HR22-00402), an ICREA professorship to F.S., the SGR funding of the Catalan government, and the Severo Ochoa centers of excellence award of the Spanish government to the hosting institution. Work in the Lehner lab was funded by European Research Council (ERC) Advanced (883742) and Consolidator (616434) grants, the Spanish Ministry of Science and Innovation (BFU2017-89488-P, EMBL Partnership, Severo Ochoa Centre of Excellence), the Bettencourt Schueller Foundation, the AXA Research Fund, Agencia de Gestio d’Ajuts Universitaris i de Recerca (AGAUR, 2017 SGR 1322), and the CERCA Program/Generalitat de Catalunya.

We thank the four members of the Flow Cytometry CRG Core Unit Erika Ramírez, Alexandre Bote, Eva Julià and Òscar Fornàs for their support and time together with Guillermo Palou for his help in retrieving the transcript sequences from the Ensembl database, and all members of Supek and Lehner labs for helpful discussions and suggestions.

## Figure legends

**Extended Data Fig. 1:**
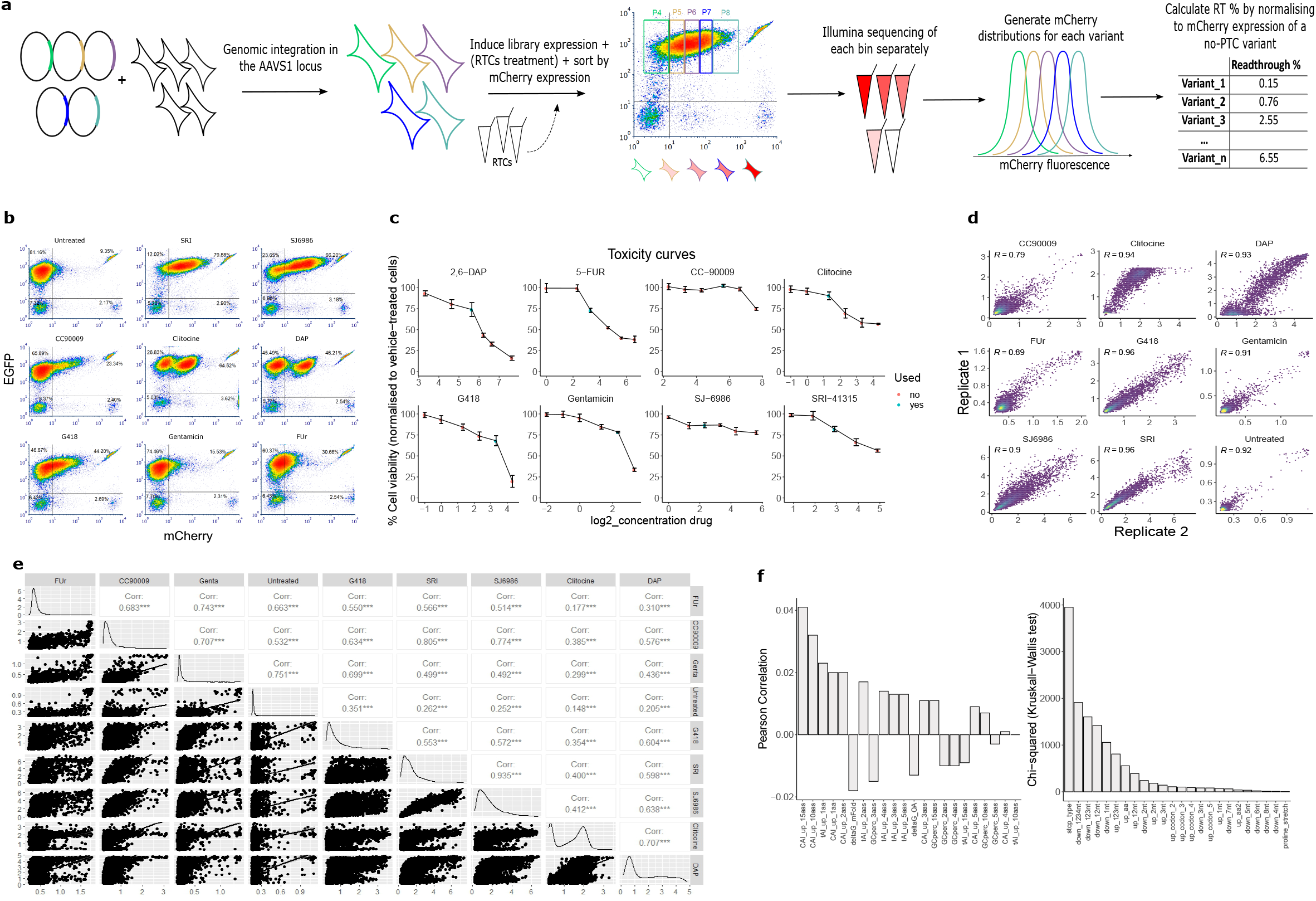
Experimental setup and overview of the drug’s datasets. **a**, Sort-sequencing overview. Each cell integrates one copy of one variant, cells are sorted based on mCherry fluorescence, bins are sequenced, and RT % are calculated from the distribution of reads of each variant normalized to the distribution of a non-PTC variant. **b,** FACS profiles (FACSAria II instrument) of the PTC library under the different treatments sorted by EGFP (Y-axis) and mCherry (X-axis). **c,** Cell viability curves for each drug. Either five or six drug concentrations were tested for each drug and the concentration displaying the highest readthrough and not reducing cell viability more than 25% (blue) was used for the assay. **d,** Inter-replicate correlations for the nine treatments. **e,** All pairwise inter-drug correlations. **f,** Sequence features associating with readthrough efficiency: showing Pearson correlations (continuous variables) and Kruskal-Wallis Chi-squared statistic (discrete variables).

**Extended Data Fig. 2:**
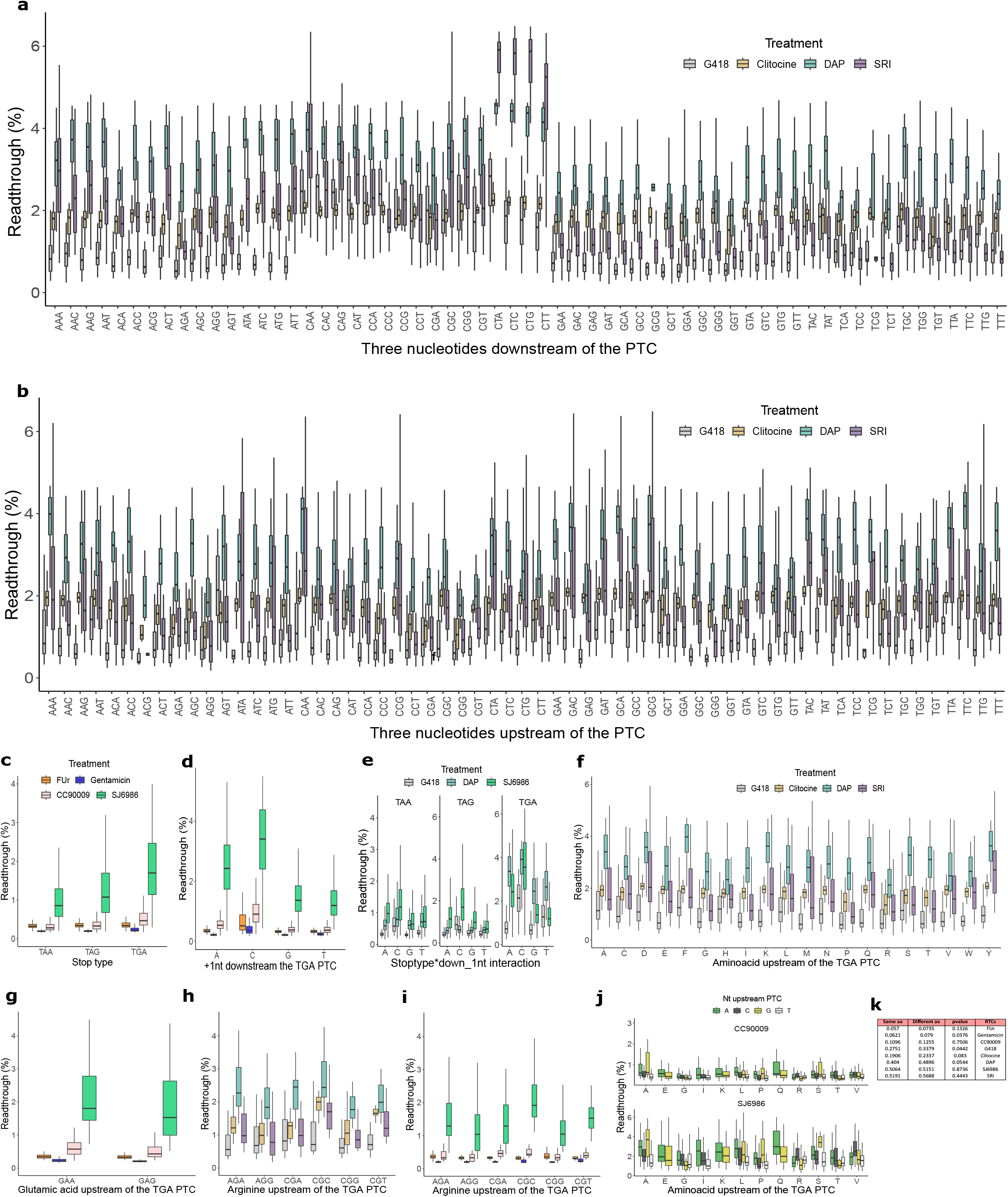
Sequence features explain the readthrough variability across PTCs and drugs. **a-i**, Effect of the sequence feature (x-axis) on readthrough efficiency (y-axis) colored by drug. The top and bottom sides of the box are the lower and upper quartiles. The box covers the interquartile interval, where 50% of the data is found. The horizontal line that splits the box in two is the median. Only variants where the stop codon is TGA are shown (except for **e**, where all stop codon variants are shown). The sequence features are the three nucleotides downstream of the PTC **(a)**, the three nucleotides upstream of the PTC **(b)**, the stop type **(c)**, the nucleotide in position +1 downstream of the PTC **(d)**, same as **d** but stratified by stop codon **(e)**, the aminoacid upstream of the PTC **(f)**, variants with a glutamic acid upstream of the PTC stratified by the codon **(g),** variants with an arginine upstream of the PTC stratified by the codon **(h, i)**. **j,** Effect of aminoacids encoded by A-ending codons on readthrough efficiency across drugs. The nucleotide in position +3 of the codon is denoted with colors. **k,** Mean readthrough difference between pairs of codons with Hamming distance of 1 (i.e. single nucleotide difference) that encode for the same amino acid, or pairs that encode a different amino acid across drugs. p-value of the t-test between same aminoacid and different aminoacid groups is shown.

**Extended Data Fig. 3:**
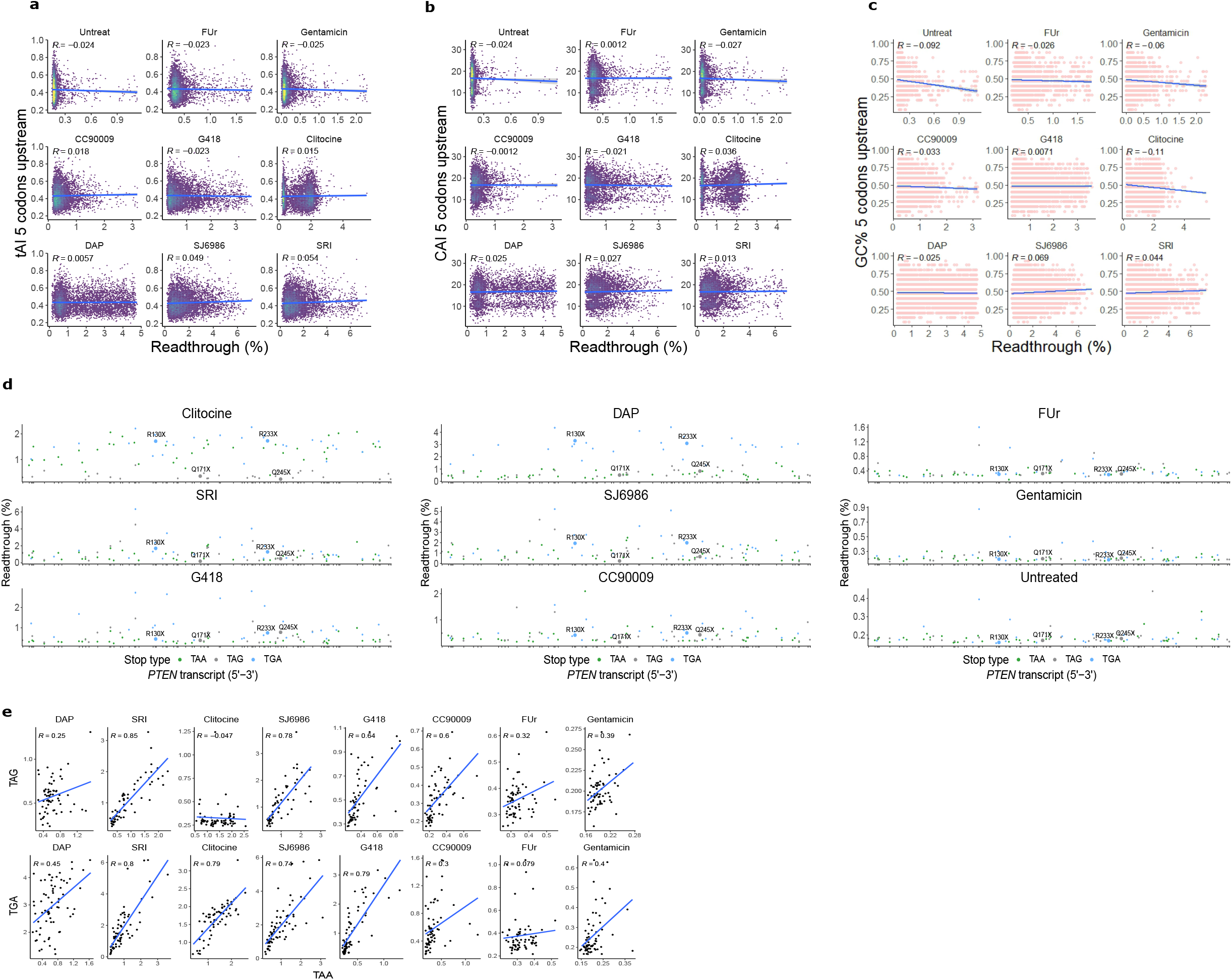
Codon related features, multistop variants and overview of PTEN nonsense mutations. **a-c**, Correlation of tAI **(a)**, CAI (**b)** and GC% **(c)** of the 5aas upstream of the PTC with readthrough efficiency for each drug. **d,** Readthrough efficiency, across drugs, for 97 nonsense *PTEN* mutations colored by stop codon type. The top-5 most recurrent nonsense mutations in the human tumor genomes are highlighted. **e,** Correlation of multistop variants across drugs. Each datapoint belongs to a mutation in the same genomic position but with a different stop type.

**Extended Data Fig. 4:**
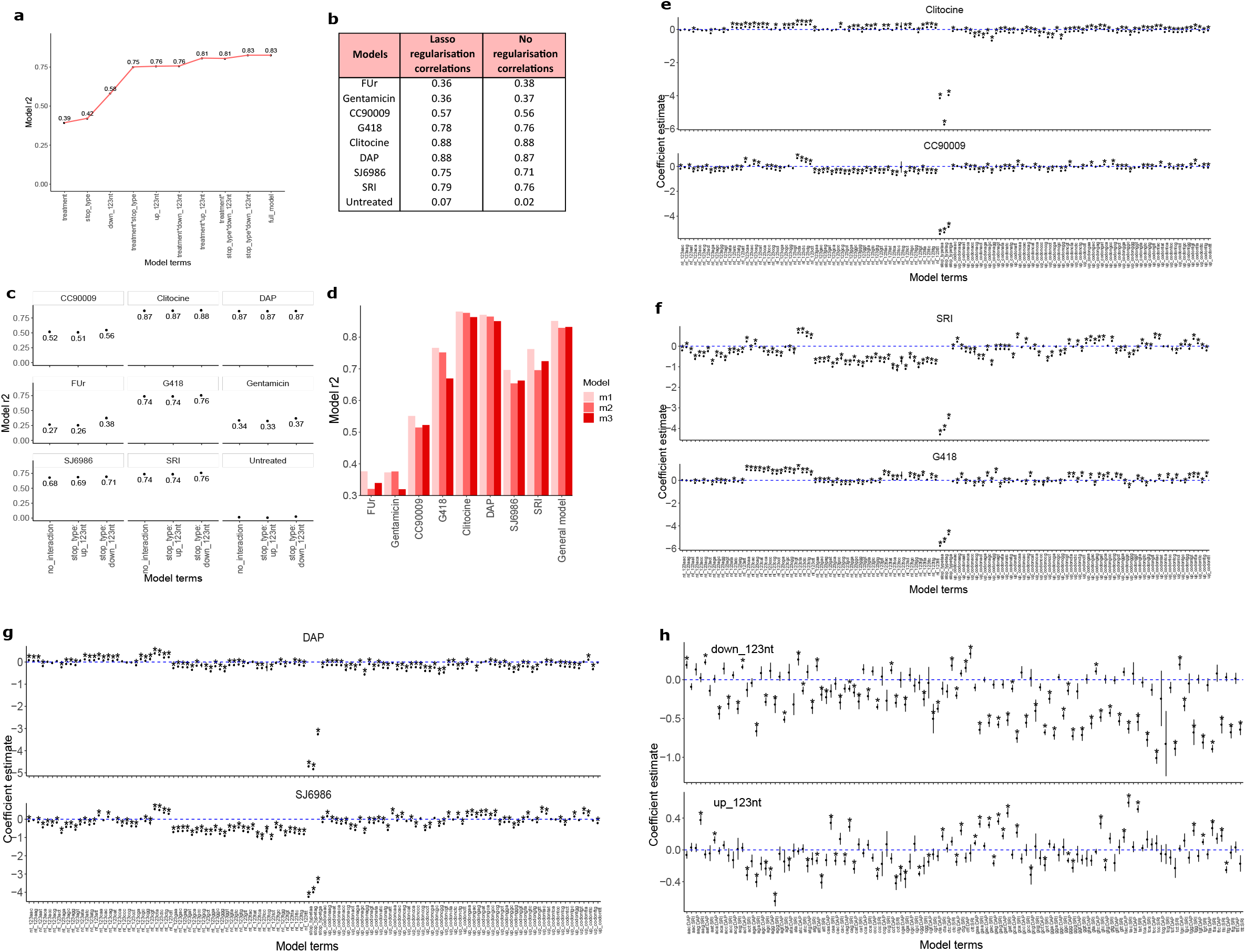
Predictive models overview and optimization. **a-i**, ANOVA test on the pan-drug model. Y-axis shows the drop in model performance upon removal of each predictive variable (X-axis). **b,** Comparison of the r^2^ for the drug-specific models when using stop type, the three nucleotides downstream and upstream of the PTC and the interaction between stop type and three nucleotides downstream versus when using Lasso regularization on 47 sequence features (Extended Data Fig.1f, Extended Data Table.6). **c,** Performance for three different model formulations across drugs: using only stop type and the three nucleotides downstream and upstream of the PTC, adding the stop type and three nucleotides upstream interaction or adding the stop type and three nucleotides downstream interaction. Only the latter consistently improves model performance across drugs. **d,** Performance for three different model formulations across drugs: encoding the three nucleotides downstream and upstream of the PTC as a nucleotide triplet (m1), encoding the upstream sequence as a nucleotide triplet and the three nucleotides downstream as three different terms (one for each position, m2) and coding the downstream sequence as a nucleotide triplet and the three nucleotides upstream as three different terms (one for each position, m3). m1 consistently higher r^2^ across drugs. **e-h,** Model coefficients for each of the drugs-specific models; Clitocine and CC90009 **(e)**, SRI and G418 **(f)**, DAP and SJ6986 **(g)** and the down_123nt (top) and up_123nt (bottom) coefficients for the pan-drug model **(h)**. Mean, 95% confidence intervals and significance (two-sided Student’s t-test) of the coefficient estimates across 10 cross validation rounds are shown. Stars represent an adjusted p-value<0.01.

**Extended Data Fig. 5:**
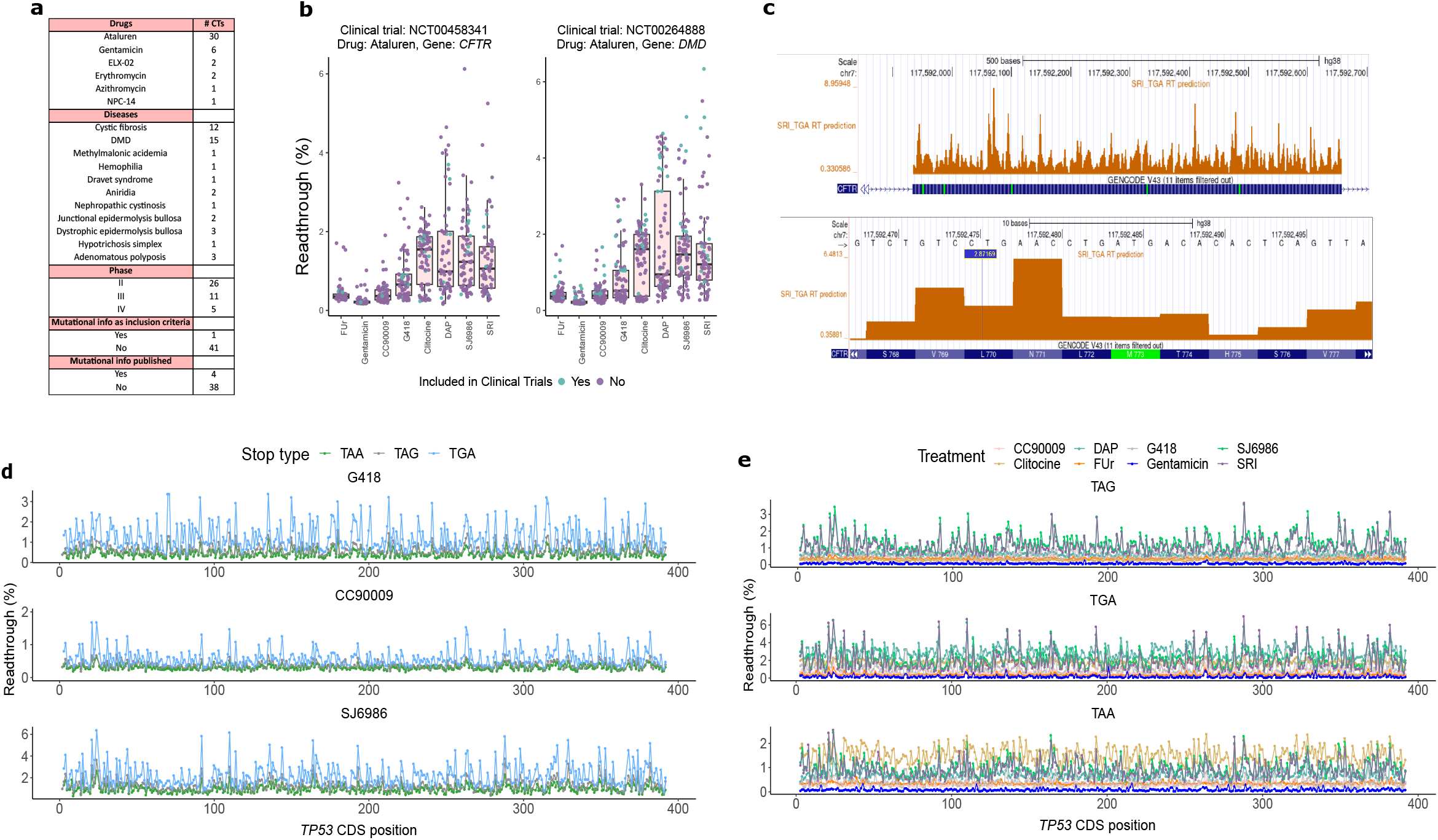
Clinical trials and *in silico* PTC saturation mutagenesis. **a**, All current (n=42) phase II-IV clinical trials testing readthrough drugs, obtained from Spelier.et.al. [^40^]. **b,** Our readthrough efficiencies of the nonsense variants in two clinical trials where genetic data was supplied (blue), together with the rest of nonsense variants in the same gene (purple). Clinical trial identifier, drug (ataluren) and gene tested are specified in the titles. Many variants included in clinical trials are unresponsive to drugs, hindering their validation. The top and bottom sides of the box are the lower and upper quartiles. The box covers the interquartile interval, where 50% of the data is found. The horizontal line that splits the box in two is the median. **c,** UCSC browser visualization of our readthrough predictions. In this specific example, SRI predictions for TGA variants are shown **d,** Overview of the readthrough predictions along the coding sequence of *TP53* for each stop codon type. Each row represents a drug-specific prediction for G418 (top), CC90009 (middle) and SJ6986 (bottom). **e,** Same as **d**, but drugs are represented as colors and each row belongs to each of the three stop types.

